# Improved eukaryotic detection compatible with large-scale automated analysis of metagenomes

**DOI:** 10.1101/2022.03.09.483664

**Authors:** Wojtek Bazant, Ann S. Blevins, Kathryn Crouch, Daniel P. Beiting

**Author notes:** Indicates co-senior authors (Kathryn Crouch, Daniel P. Beiting).

## Abstract

**Background:** Eukaryotes such as fungi and protists frequently accompany bacteria and archaea in microbial communities. Unfortunately, their presence is difficult to study with ‘shotgun’ metagenomic sequencing since prokaryotic signals dominate in most environments. Recent methods for eukaryotic detection use eukaryote-specific marker genes, but they do not incorporate strategies to handle the presence of eukaryotes that are not represented in the reference marker gene set, and they are not compatible with web-based tools for downstream analysis.

**Results:** Here we present CORRAL (for Clustering□Of□Related Reference ALignments), a tool for identification of eukaryotes in shotgun metagenomic data based on alignments to eukaryote-specific marker genes and Markov clustering. Using a combination of simulated datasets, mock community standards, and large publicly available human microbiome studies, we demonstrate that our method is not only sensitive and accurate but is also capable of inferring the presence of eukaryotes not included in the marker gene reference, such as novel strains. Finally, we deploy CORRAL on our MicrobiomeDB.org resource, producing an atlas of eukaryotes present in various environments of the human body and linking their presence to study covariates.

**Conclusions:** CORRAL allows eukaryotic detection to be automated and carried out at scale. Implementation of CORRAL in MicrobiomeDB.org creates a running atlas of microbial eukaryotes in metagenomic studies. Since our approach is independent of the reference used, it may be applicable to other contexts where shotgun metagenomic reads are matched against redundant but non-exhaustive databases, such as identification of bacterial virulence genes or taxonomic classification of viral reads.

## Background

Eukaryotic microbes are a large and phylogenetically diverse group of organisms that includes both pathogens and commensals, the latter of which are emerging as important modulators of health and disease. Protists include many important pathogens of humans and other animals, such as *Cryptosporidium, Toxoplasma, Eimeria, Trypanosoma*, and *Plasmodium*. Many fungi are also well-studied pathogens affecting a diverse range of hosts. For example, *Aspergillus fumigatus* is an important cause of respiratory disease in humans (1); *Magnaporthe oryzae* is the most important fungal disease of rice globally (2); while *Pseudogymnoascus destructans* is the cause of White-Nose Syndrome, one of the most devastating diseases of bats (3). However, recent data also suggest that non-pathogenic commensal fungi are critical modulators of the human antibody repertoire (4–6), intestinal barrier integrity (7), and colonization resistance (8). The diverse array of host-microbe interactions and host phenotypes influenced by eukaryotic microbes underscores the importance of studying this class of organisms in their natural habitats. Unfortunately, the ability to carry out culture-independent analysis of eukaryotic microbes is severely hindered by their low abundance relative to bacteria, which makes accurate detection a challenge and consequently eukaryotes are commonly overlooked in metagenomic studies (9). For example, an analysis of stool metagenomes in healthy adults participating in the Human Microbiome Project reports only 0.01% reads aligning to fungal genomes (10).

Several methods have been developed to improve the detection of eukaryotes in complex samples. Targeted sequencing of internal transcribed spacer regions (ITS) is a common approach but prevents simultaneous profiling of other members of the microbiome (11). Alternatively, collections of curated fungal genomes have been successfully used for strain-level identification of *Blastocystis* from stool (12). However, pitfalls associated with non-specific or erroneous parts of reference genomes (13) combined with computational challenges associated with carrying out alignments to very large collections of reference genomes (14) limit applicability of these approaches to the discovery of eukaryotes from the vast amount of metagenomic data already available in the public domain. One attractive solution to this challenge was recently proposed in important work by Lind and Pollard (15), who base their method for sensitive and specific identification of eukaryotes in metagenomic studies, EukDetect, on alignments to a collection of over 500,000 universal, single-copy eukaryotic marker genes.

We recently sought to add the EukDetect reference database and software to our web-based resource, MicrobiomeDB.org (16), to allow for automated detection of eukaryotes across a range of human metagenomic studies currently available on the site. Since the EukDetect pipeline does not allow for adjustment of filtering thresholds and is not packaged for containerized deployments, we decided to implement our own tool built with a more flexible software architecture. Our approach retains the EukDetect reference database, as well as the use of Bowtie2 (17) since it has been shown to be a sensitive aligner (18). To better understand the filtering process used by EukDetect, we carried out a simulation-based evaluation. We observed that filtering of read alignments based on mapping quality (MAPQ) scores (19) – though necessary for EukDetect’s high specificity – removes correct alignments for which Bowtie2 has inferior but closely scored alternatives.

Considering that the difficulty of detecting a taxon may be affected by similarity of its marker gene sequences to its most similar neighbor led us to develop CORRAL (for Clustering□Of□Related Reference ALignments), an approach for processing marker gene alignments based on exploiting information in shared alignments to reference genes through Markov clustering. This allows for sensitive and accurate detection which also extends to species not present in the reference, but which are similar to one or more known taxa present in the reference.

## Results

### Species-specific impact of MAPQ filtering

EukDetect relies on mapping quality (MAPQ) scores (19) for its performance. To evaluate how read mapping and filtering parameters influence eukaryotic detection, we carried out a series of simulations using the EukDetect database of marker genes as both a source of reads with known identity and a reference to which to align these reads. When metagenomic reads are simulated from this reference and then simply mapped back, thus exactly matching the reference, they are accurately mapped to the correct taxon with a recall (fraction of correctly mapped reads among all reads) and precision (fraction of correctly mapped reads among all reads that mapped) of 95.1% for each. Applying a MAPQ ≥ 30 filter increases precision to 99.7% and decreases recall to 91.7%. This translates to 92% of the simulated reads mapping with MAPQ ≥ 30, with only 0.3% of these mapping incorrectly, and out of the remaining 8%, almost half mapping incorrectly.

Examining these data at the level of individual taxa from which the reads were sourced reveals a structural component to the difficulty of mapping the reads, as well as the efficacy of the MAPQ filter (**Figure 1**). For example, out of 3977 taxa whose reads map back to the reference, reads from 1908 taxa map with 100% precision (**Figure 1, upper rightmost points**), and after applying the MAPQ ≥ 30 filter, 1105 more taxa map with 100% precision. Despite this clear improvement after filtering, 146 taxa still map with precision lower than the pre-filter overall total of 95.1% (**Figure 1, dashed line**). This set of taxa includes numerous species of *Aspergillus* (**Figure 1A**), *Leishmania* (**Figure 1B**), and *Trichinella* (**Figure 1C**), all of which are important pathogens of humans and other mammals. Furthermore, filtering based on MAPQ decreases precision for five taxa, including the fungi *Fusarium cf. fujikuroi NRRL 66890* and *Escovopsis sp. Ae733* (**Figure 1A**), and the protists *Favella ehrenbergii, Leishmania peruviana*, and *Mesodinium rubrum* (**Figure 1B**). Taken together, these results suggest that relying on MAPQ filter alone may not allow for robust detection of multiple eukaryotes of public health importance.

**Figure 1:**
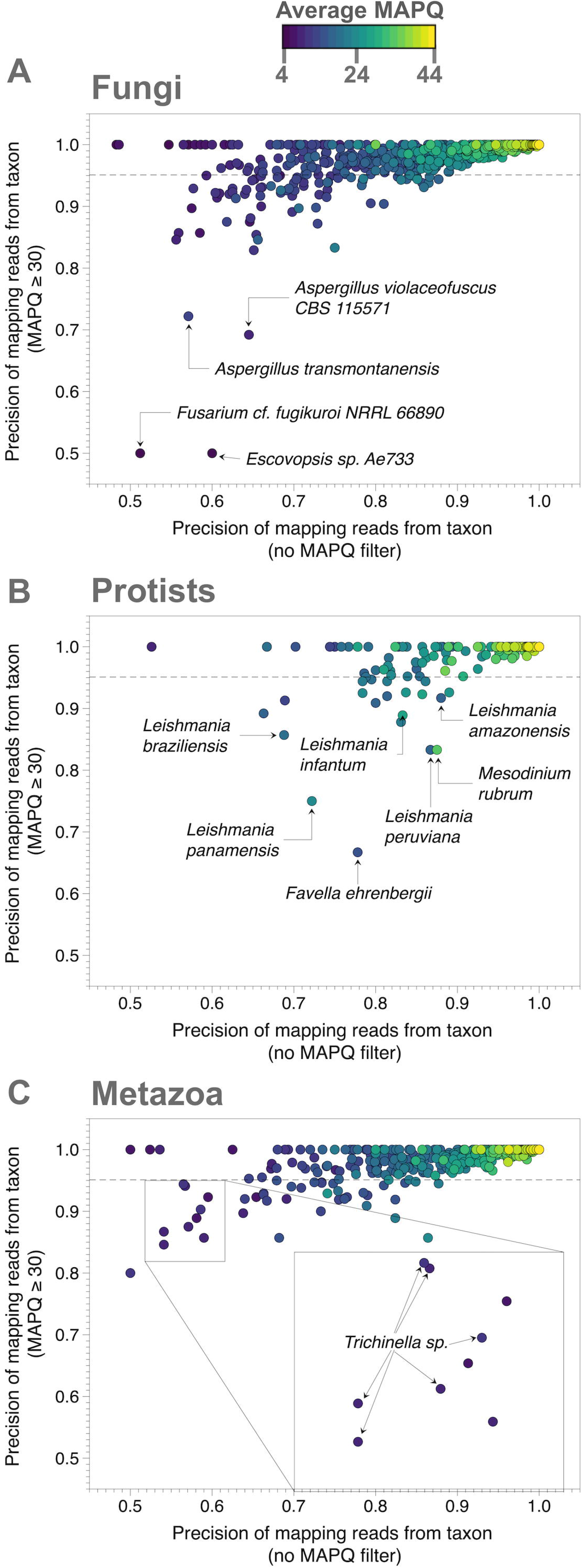
Species-specific impact of MAPQ filtering. Precision of read mapping comparing MAPQ ≥ 30 (Y-axis) versus no MAPQ filter (X-axis) for A) fungi, B) protists, and C) metazoa. Points are colored by average MAPQ scores. Horizontal dashed line indicates prefilter precision and recall of 95.1%. Select taxa for which the MAPQ filter either only marginally improved or impaired precision are labeled.

Since the diversity of eukaryotic microbial life extends far beyond the currently discovered species, let alone species present in the EukDetect reference (20), we next modified the simulation above to evaluate the possibility of detecting ‘unrepresented’ species. To do this, species-level markers in the EukDetect reference were split into a holdout set of 371 taxa from which we simulated reads that were then mapped back to the remaining 3343 taxa in the EukDetect reference, thus mimicking a scenario in which a metagenomic sample contains reads from eukaryotes not represented in the reference. In this circumstance, the MAPQ ≥ 30 filter is not on average an improvement. Same-genus precision and recall are 82% and 30%, respectively, without the filter. Applying the MAPQ filter results in a similar precision (83.6%) but a much-diminished recall of 7%. Source taxon is a structural component here as well – applying the MAPQ ≥ 30 filter increases the number of taxa which only map to the correct genus from 48 to 152 but increases the number of taxa that fail to map from 49 to 175.

There is extensive strain variation in complex microbial communities, so we next set out to evaluate the ability to identify eukaryotes when a sample contains a novel strain of a species present in the reference database **(Figure 2)**. We carried out a third simulation in which sampled reads were mutated before mapping back to the reference. As mutation rate increases, recall declines from 95.1% to less than 10% when mutation rate is 0.2. In this range, precision stays between 95-96% for all reads and ≥ 99% for reads with MAPQ ≥ 30 – an observation consistent with previous reports of bowtie2 preserving precision over recall [22]. Applying the MAPQ ≥ 30 filter results in a rapid decline in recall. For example, when mutation rate is 0.1, recall is 68.3% overall but drops to 5.0% when a MAPQ ≥ 30 filter is applied. These results indicate there may be many taxa which match the reference sufficiently closely to allow for sensitive detection, but only if one does not apply the MAPQ ≥ 30 filter.

**Figure 2:**
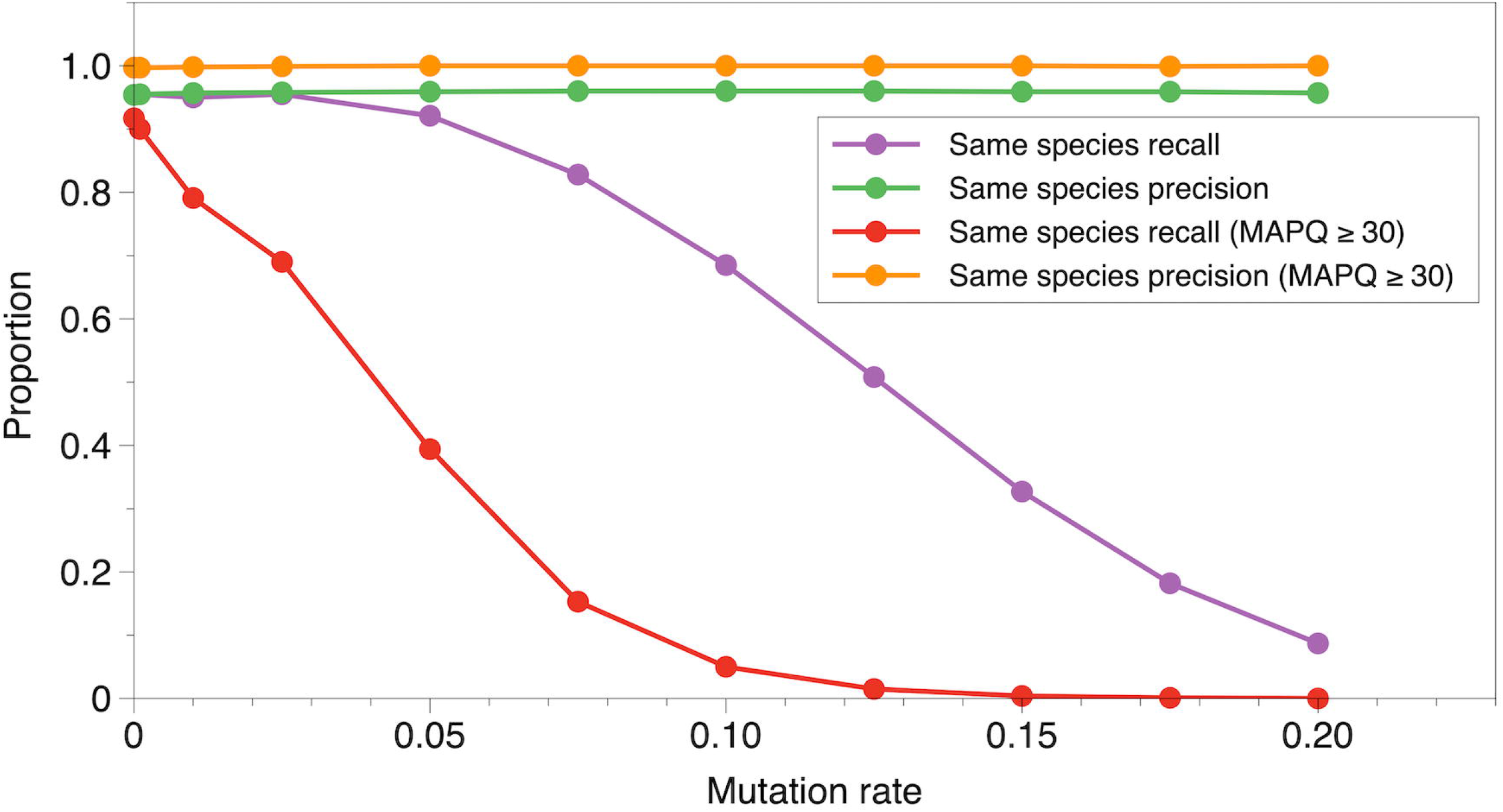
Mutation rate influences MAPQ filter performance. Proportion of taxa where recall or precision are as described (legend), as mutation rate is increased from 0 to 0.2.

### CORRAL leverages Markov clustering for reference-based eukaryote detection

To address the challenges described above and to fully leverage the valuable eukaryotic marker gene reference database created by Lind and Pollard (15), we developed CORRAL (Clustering□Of□Related Reference Alignments) as a Nextflow workflow wrapping a Python module. CORRAL retrieves sequence files, aligns reads to the EukDetect reference of markers, and produces a taxonomic profile through a multi-step process (**Figure 3**). First, we run Bowtie2 and keep all alignments that are at least 60 nucleotides in length (**Figure 3, step 1**), ensuring that sequence matches contain enough information to be marker specific. We then run Markov Clustering (MCL) on a graph composed of marker genes as nodes and counts of shared alignments as edge weights to obtain marker clusters (**Figure 3, step 2**). Next, percent match identities of alignments are calculated and aggregated by marker to obtain an identity average for each marker gene, as well as per cluster to obtain a cluster average (**Figure 3, step 3**). Each marker whose identity average is lower than the cluster average is considered an inferior representation of signal in the sample, and taxa with ≥ 50% of such markers are rejected (**Figure 3, step 4**). Remaining taxa are then gathered into taxonomic clusters using MCL on counts of multiply aligned reads (**Figure 3, step 5**), which allows us to incorporate ambiguity of identification into any taxa reported. Unambiguous matches (defined as having average alignment identity of ≥ 97%, and two different reads aligned to at least two markers) are reported (**Figure 3, step 6**), while other taxa in clusters where there are any unambiguous matches reported are rejected. Finally, for each remaining taxon cluster, we report it as one hit if it is a strong ambiguous match (defined as having at least four markers and eight reads) by joining names of taxa in the cluster and prepending with a “?” (**Figure 3, step 7**). See methods for a description of threshold selection.

**Figure 3:**
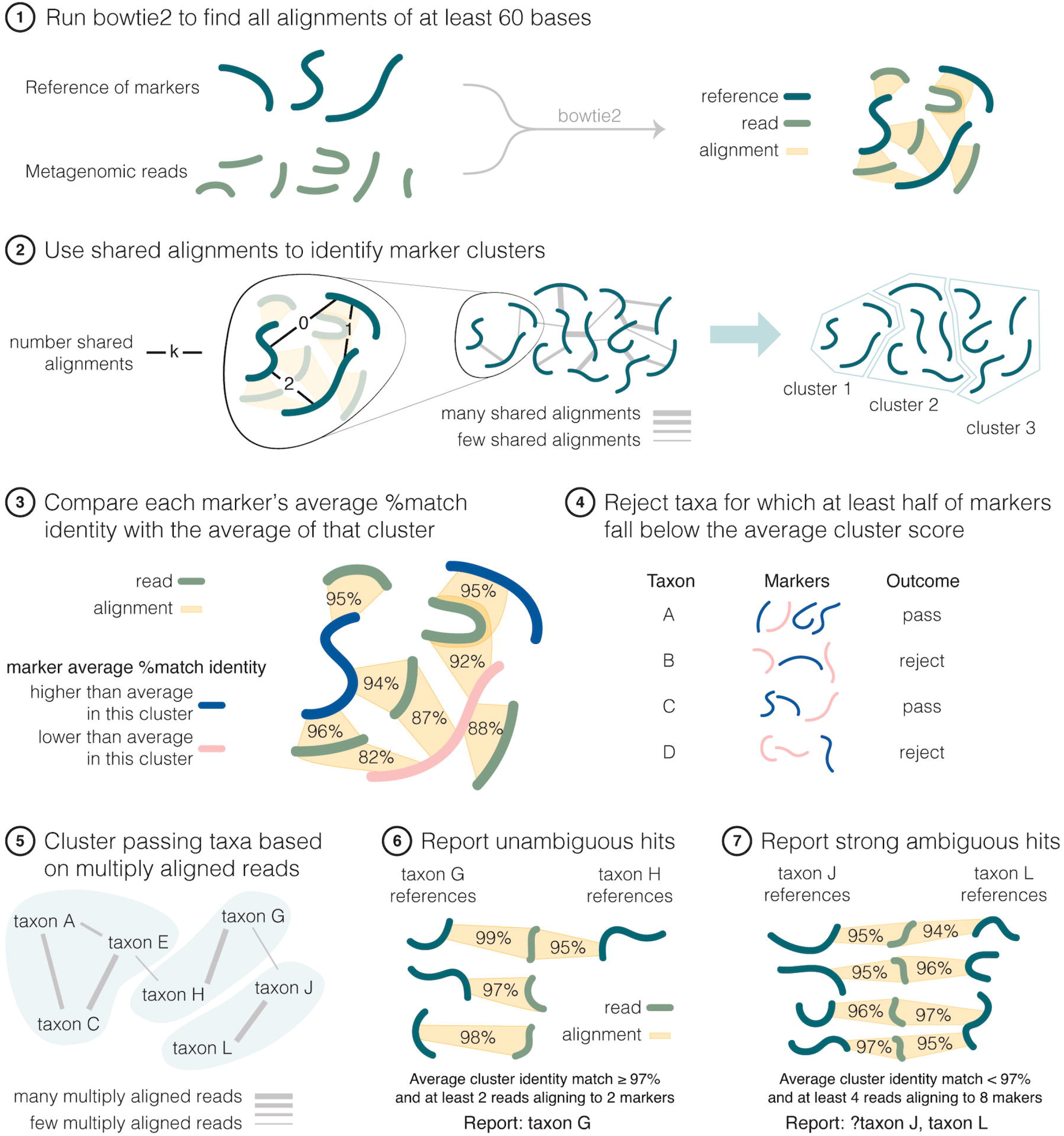
The CORRAL workflow. Schematic showing all seven steps of the CORRAL workflow.

This approach represents a set of default parameters – based on our observations in simulated and human microbiome data – that can be altered when configuring CORRAL. Additionally, CORRAL has rich reporting capabilities, including the ability to quantify abundance of eukaryotes using a ‘copies per million (CPM)’ metric (see Methods).

### CORRAL detects low abundance taxa and reports unrepresented species

Microbial eukaryotes are often present at low abundance in metagenomic studies, underscoring the importance of evaluating the performance of detection software at low limits of detection. By design, CORRAL can detect a species when as few as two reads each align to a different reference marker. For EukDetect the lower limit of detection is four reads (or two paired-end reads). To systematically evaluate the performance of both tools, we prepared 338 simulated samples – each containing a single taxon at this minimal abundance – and determined the sensitivity and specificity of CORRAL compared to EukDetect with either default or sensitive settings. A tool with perfect sensitivity and specificity would detect the low abundance taxon present in each of the 338 simulated samples, with no additional taxa reported. Using only Bowtie2 for read mapping, without additional specialized eukaryotic detection software, sensitivity was 100% and specificity was 93.4% **(Figure 4A, red bars)**. EukDetect with default settings resulted in only 72.5% sensitivity, but perfect specificity **(Figure 4A, blue bars)**. Using either EukDetect in sensitive mode or CORRAL, sensitivity and specificity were similar and above 95% **(Figure 4A, green and purple bars, respectively)**. These results suggest that correctly identifying a eukaryote that exactly matches the reference – even if only present at the lower limit of detection – does not require use of a MAPQ filter or Markov clustering, but that both of these improve specificity.

**Figure 4:**
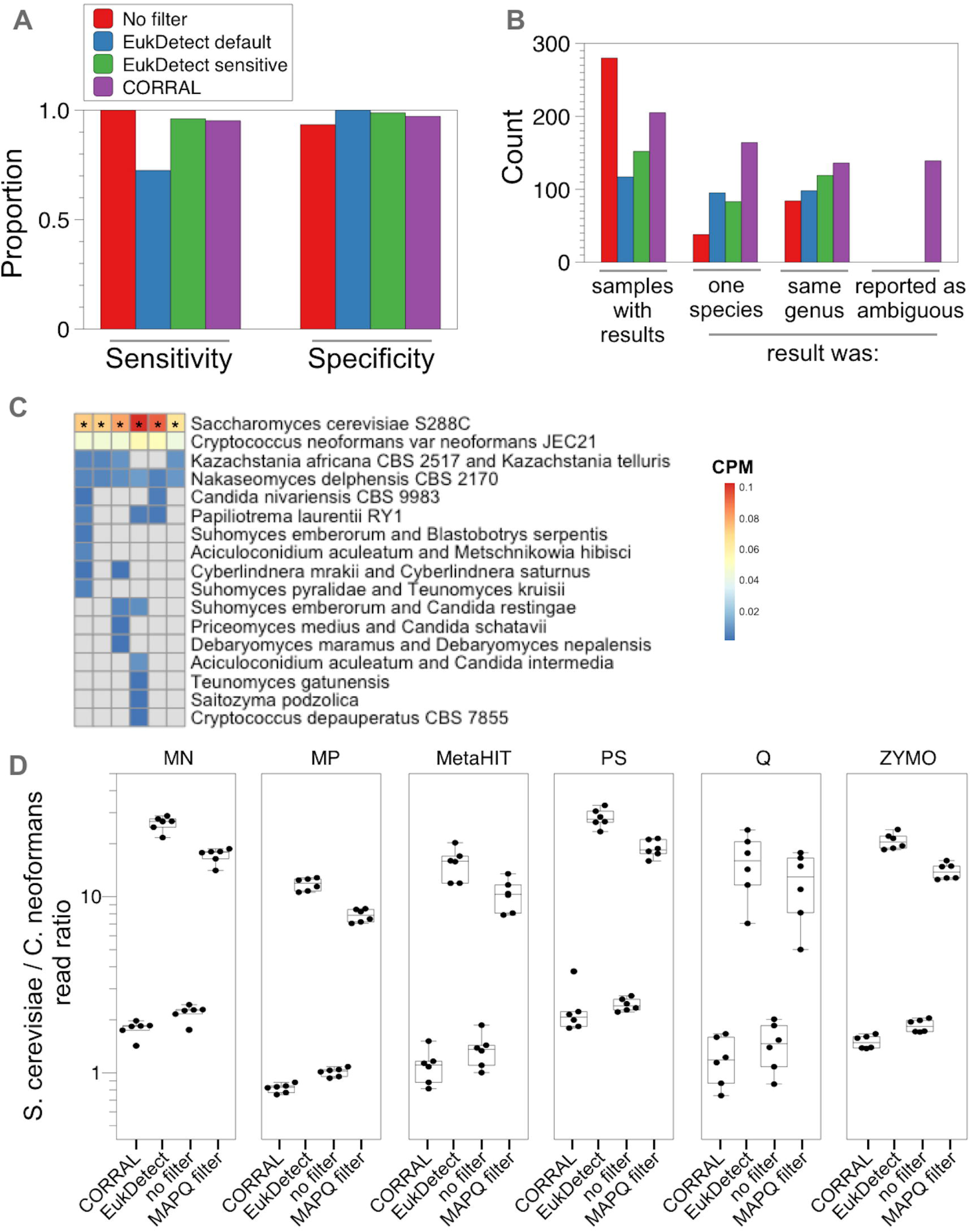
CORRAL yields high sensitivity and specificity when predicting the presence of eukaryotes in metagenomic data. A) Proportion of results (Y-axis) yielding the correct species at lower limit of detection for EukDetect and CORRAL. B) Unrepresented species simulation results. Number of samples (count; Y-axis) are shown for which results were produced by EukDetect or CORRAL (X-axis). C-D) ZymoBIOMICS mock community standard from Yang et al. (21). C) heatmap showing copies per million (CPM) for all microbial eukaryotes detected by CORRAL in 6 replicate samples of the ZymoBIOMICS standard extracted using the zymo protocol as described previously (21). Asterisks mark unambiguous results from CORRAL. D) Plots showing ratio of reads assigned to *S. cerevisiae* versus *C. neoformans* for CORRAL, EukDetect, ‘no filter’ and ‘MAPQ filter’ (see methods). Separate box plots are shown for each extraction method used, including MagPure Fast Stool DNA KF Kit B (‘MP’); Macherey Nagel NucleoSpin Soil kit (‘MN’); Zymo Research Quick-DNA Fecal/Soil Microbe kit (‘ZYMO’); protocol Q (‘Q’); MOBIO DNeasy PowerSoil kit (‘PS’); and a non–kit-based manual protocol adopted by the Metagenomics of the Human Intestinal Tract Consortium (‘MetaHIT’) Each point represents a separate mock community sample that was extracted, sequenced, and analyzed.

The EukDetect marker gene collection, like all microbial marker gene references, is incomplete. Thus, we next evaluated how these software tools handle reads from a taxon that is not provided in the reference. We returned to our holdout simulation described above and simulated a metagenomic dataset consisting of 338 samples, each containing a single ‘unrepresented’ eukaryotic species (from the holdout set) at 0.1x genome coverage **(Figure 4B)**. Using this data set, we again compared CORRAL, EukDetect default, and EukDetect sensitive, this time by whether a single species was reported by each tool, and whether the reported species belonged to the same genus as the unrepresented taxon from the holdout. Of the three methods tested, CORRAL returned the most samples with results (205/338; **Figure 4B, purple bar**), the most results with one species reported (164/338), and the most results in the same genus (136/338). Importantly, since the unrepresented taxon in each sample, by definition, is not present in the reference, ideally a tool should report some level of uncertainty for each sample. CORRAL, but not EukDetect, can report strong but ambiguous results that do not match perfectly to the reference **(Figure 3, step 7)**, and did so for 139 out of the 205 samples for which it returned any results **(Figure 4B)**. Collectively, these results highlight that CORRAL is a sensitive method to detect microbial eukaryotes but is also capable of reporting uncertainty in results, thus empowering users to interpret results more easily.

We next set out to evaluate the performance of CORRAL and EukDetect on samples where ground truth is known. We analyzed publicly available data (21) from the ZymoBIOMICS mock community standard which contains 8 bacterial species and two fungal species (*Saccharomyces cerevisiae* and *Cryptococcus neoformans*). Yang et al. extracted DNA from this community standard using six different methods, in order to assess the extent to which community composition is impacted by extraction method (21). Analysis of this data by CORRAL identified both *S. cerevisiae* and *C. neoformans*, but the latter was flagged by CORRAL as ambiguous **(Figure 4C)**. This ambiguity is likely due to the strain used in the mock community being different from the *C. neoformans* strain present in the marker gene reference, as evidenced by reduced number of reads aligning and lower % identity, compared to *S. cerevisiae*, for both EukDetect and CORRAL **(Table S1)**. CORRAL also yielded several false positive taxa, all of which were flagged as ambiguous **(Figure 4C)**. In contrast, EukDetect yielded perfect sensitivity and specificity for both fungal taxa across all extraction methods **(Table S1)**. Both *S. cerevisiae* and *C. neoformans* are present in the community standard at a theoretical relative abundance of 2%. Consistent with this notion, the ratio of reads assigned by CORRAL to *S. cerevisiae* versus *C. neoformans* was between 1 and 2 – depending on the extraction method used – indicating that CORRAL accurately estimated roughly similar relative abundance for these two taxa **(Figure 4D)**. In contrast, EukDetect predicted a relative abundance of *S. cerevisiae* that was at least an order of magnitude higher than that of *C. neoformans*, independent of the extraction method used **(Figure 4D)**. To test whether the MAPQ filter was source of the difficulty in estimating abundance by EukDetect, we analyzed the community standards using only the best alignment and a simple read length filter **(Figure 4D, ‘no filter’)**. As expected, this resulted in high sensitivity but extremely poor specificity **(Table S1)** and yielded relatively balanced abundance estimates for both fungal taxa **(Figure 4D)**. In contrast, adding the MAPQ filter resulted in abundance estimates that were dramatically skewed in favor of *S. cerevisiae*. Taken together these data suggest that CORRAL balances high sensitivity for detection with accurate estimation of relative abundance, while also reporting uncertainty.

### Understanding the impact of species relatedness on microbial eukaryote detection

Metagenomic samples are complex and may contain closely related species, which could impact the sensitivity or specificity of a eukaryotic detection tool. We reasoned that the extent to which detection software is affected by potentially ‘confusable’ species likely depends on the species-species pair in question. To rigorously evaluate this, we return to the same database of alignments of simulated reads used in Figure 1 and evaluate all pairs of taxa to measure the proportion of reads sampled from a source taxon that align to a different taxon. For each member of a pair of taxa, we computed the rates at which a taxon either emits or accepts reads from either the other member of the pair, or from other taxa outside of the pair **(Figure S1A)**. This resulted in 8 rate computations for each of 4558 pairs of taxa **(Figure S1B; Table S2)**. Principal component analysis (PCA) of this data yielded a reduced dimensional space where each point represents a single pair of taxa **(Figure S1C)**. PC1 accounted for 61.6% of the variance, its coefficients were positive for all features, and were symmetric between the pair members, so we interpret it as a representation of the extent to which a pair of taxa were easily confused with each other or with taxa outside of the pair. PC2 explained 21.5% of the variance and represented bias in the ability to identify one versus the other member of the pair. Pairs positioned high or low on PC2 were more difficult to correctly identify taxon A or B, respectively **(Figure S1C)**. This relatively simple mathematical representation of the pairwise ‘confusability’ of taxa in metagenomic samples provides a useful framework to evaluate any eukaryotic prediction tool.

We next examined this confusability space in more detail to understand whether taxonomy played a role in the potential difficulty of detecting members of a given pair **(Figure 5A)**. Notably, most points were concentrated in a low confusability area that is positioned low on PC1 and centered on PC2 **(Figure 5A and 5B)**, suggesting that most microbial eukaryote pairs should be easily distinguished from each other and from other taxa outside of the pair, irrespective of the software used. Interestingly, this area of the PCA plot contains the pairing of *Entamoeba histolytica* and *Entamoeba dispar* **(Figure 5A, arrow)**, the former an important gastrointestinal parasite and the latter a harmless gut commensal. This pair was highlighted by Lind and Pollard as an example of the utility of the EukDetect software (15). Our data show that the difficulty in distinguishing related pairs depends on the genus in question. For example, many pairings of *Aspergillus* species showed low confusability. In contrast, *Leishmania* and *Fusarium* species both showed intermediate difficulty, while species of the parasitic helminth, *Trichinella*, showed high confusability **(Figure 5A and 5C)**.

**Figure 5:**
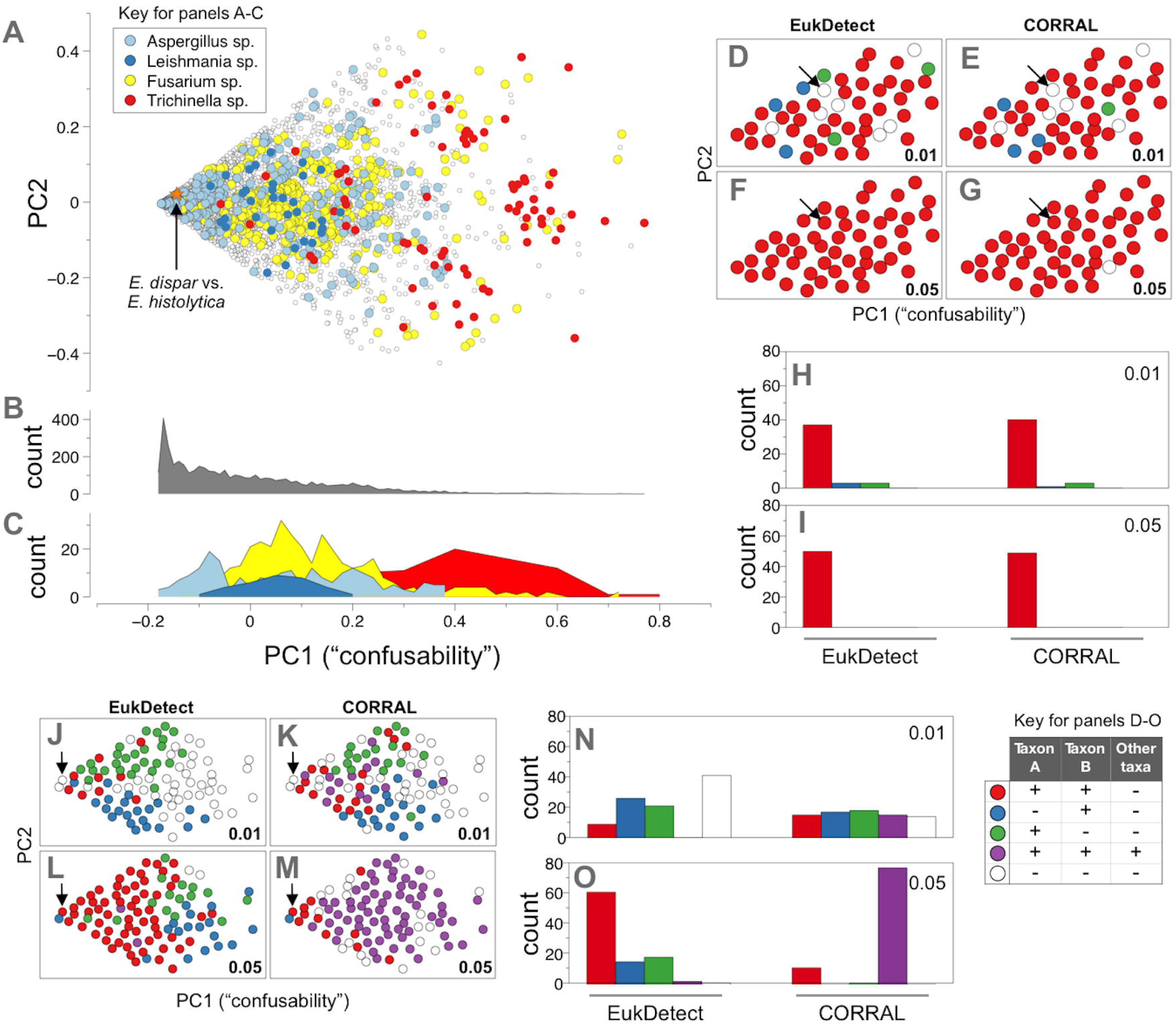
Identifying closely related eukaryotes in metagenomic samples. (A) Principal component analysis showing the confusability for 4558 pairs of microbial eukarote species. Each point represents a pair of taxa. Colors indicate pairs where both members belong to the genera of *Aspergillus* (light blue), Leishmania (dark blue), *Fusarium* (yellow) or *Trichinella* (red), selected because they differ in confusability. Arrows throughout indicate point corresponding to the *Entamoeba dispar/Entamoeba histolytica* pair. Histograms for (B) all points or (C) only points corresponding to the selected genera in the PCA plot from panel A. (D-G) Focused subset corresponding to low confusability area around the *E. dispar/ E. histolytica* pair in panel A, sampled at either low (0.01x; panels D and E) or higher coverage (0.05x, panels F and G). (H-I) bar plot summaries of panels D-G. (J-K) Broad subset biased to include samples with high confusability. (N-O) bar plot summaries of J-K. Colors for panels D-O reflect sensitivity and specificity of predicting the presence of both members of a pair.

Next, we set out to evaluate the extent to which CORRAL and EukDetect could identify pairs of taxa that spanned a wide range in potential confusability **(Figure 5A)**. We first sampled 50 pairs near the low confusability area around the *E. histolytica* and *E. dispar* pair (see Methods). Even when pairs had low coverage (0.01x), both tools were able to identify both taxon A and B for the vast majority of pairs examined (**Figure 5D, 5E, and 5H**), but both failed to correctly identify *E. histolytica* and *E. dispar* (**Figure 5D and 5E**, arrows). At higher coverage (0.05x) both tools not only identified *E. histolytica* and *E. dispar* (**Figure 5F and 5G**, arrows), but also demonstrated perfect or near perfect sensitivity and specificity for all 50 pairs **(Figure 5F, 5G, and 5I)**. To expand on this analysis and assess a more challenging set of pairs, we repeated our sampling from Figure 5A, but selected 98 pairs that spanned a broader range of confusability **(Figure 5J-O)**. Running EukDetect and CORRAL on this subset revealed the difficulty of correctly identifying pairs of taxa as confusability increases. At low coverage (0.01x) EukDetect correctly identified only 9 of the 98 pairs, while CORRAL exhibited higher sensitivity and detected 15 pairs (**Figure 5J and 5K, red points)**, but this increased sensitivity came at the expense of specificity (**Figure 5J and 5K, purple points)**. EukDetect and CORRAL are most successful at reporting both taxon A and taxon B and no other taxa for lower values of PC1 **(Figure 5J and 5K, red points)**. When reporting incomplete results, the tendency of both tools to detect either taxon A **(Figure 5J and 5K, green points)** or taxon B **(Figure 5J and 5K, blue points)** depends on the sign of PC2, reinforcing the notion that our approach **(Figure S1)** provides a confusability map of microbial eukaryote pairs. The increased sensitivity of CORRAL resulted in only 14 out of 98 pairs (14%) without results at 0.01x coverage, while EukDetect left 41 pairs (42%) without results. Finally, as coverage increases **(Figure 5L, 5M, and 5O)**, EukDetect shows high sensitivity and near perfect specificity, while CORRAL shows high sensitivity but poor specificity.

### Evaluating CORRAL on human microbiome data

To move beyond the simulations described above we next tested CORRAL on data from real microbiome studies where some expectations exist about which eukaryotes might be present. We first evaluated the DIABIMMUNE study (22), for which 136 data points about 30 different eukaryotes were reported across 1154 samples in the original EukDetect publication (15). Processing these same 1154 samples, CORRAL is in exact concordance with EukDetect on 122/136 data points and adds an additional 97 data points. CORRAL reports common taxa at a higher frequency. For example, *S. cerevisiae* is detected by CORRAL 67 times, while EukDetect only identifies this organism 31 times. The other additional hits detected by CORRAL, but not EukDetect, consist primarily of yeast and other fungi that have been previously reported in the human gut. Importantly, CORRAL differs from EukDetect in how it treats reads that might originate from a unrepresented species. For example, in sample G78909 from DIABIMMUNE, EukDetect reports *Penicillium nordicum*, while our method reports a novel *Penicillium*. In sample G80329, our method agrees with EukDetect regarding detection of *Candida parapsilosis*, while also identifying the sample as positive for *C. albicans*. Finally, in sample G78500 EukDetect reports *Saccharomyces cerevisiae* and *Kazachstania unispora*, which our method reports to be reads from a single taxon: a strain of *S. cerevisiae* that differs from the reference strain. The additional taxa detected by CORRAL seem plausible, given that they are common gut-associated fungi, but since we lack a ground truth for eukaryotes in this or any other metagenomic study we cannot know whether the results produced by CORRAL or EukDetect more accurately reflect the true microbial eukaryote community in these samples.

### Automating eukaryote detection with CORRAL

In addition to making our software simple to install through pip and easily parametrized, we integrated CORRAL into the automated data loading workflow for our open-science platform, MicrobiomeDB.org. As of Release 30 (9 Nov 2022), the site contains 6337 samples from 8 published metagenomic studies (22–29). Automated analysis of these samples by CORRAL occurs at the time a study is loaded for public release onto the database website, and microbial eukaryote data becomes readily available to users through a sophisticated web toolkit **(Figure S2)**. For example, selecting the DIABIMMUNE study on the site and navigating to ‘Microbial eukaryote analysis’ **(Figure S2, red rectangle)** reveals two ways that CORRAL data is represented on the site: detection and abundance **(Figure S2, top and bottom panel, respectively)**. Selecting ‘Fungal taxon detected by sequence match’ **(Figure S2A)** presents a multifilter that lets users view and select samples positive (‘Y’) for any fungal taxon **(Figure S2B)**. For example, users can easily find that CORRAL detected *Saccharomyces cerevisiae* in 67 samples (6%) from the DIABIMMUNE study **(Figure S2C)**. Abundance data is available under ‘Normalized number of taxon-specific sequence matches’ **(Figure S2D**). Selecting a single taxon, such as *Candida parapsilosis* **(Figure S2E)**, shows a distribution of abundance of that taxon across all samples in the study, thus making it easy to view and select samples with high levels of any taxon of interest **(Figure S2F)**.

CORRAL identified microbial eukaryotes in 1453/6337 (23%) of the metagenomic samples on MicrobiomeDB, yielding 2084 data points for 190 different eukaryotic taxa. A large majority, 1851/2084 or 89% of these data points, are fungal taxa. Of the 233 data points for non-fungal eukaryotes detected in these samples, 200 (86%) are species belonging to the genus *Blastocystis*, one of the most common protozoan parasites found in the human GI tract (30). A summary of the top 15 most frequently observed eukaryotes (**Table 1**) reveals that *Malassezia restricta*, a common commensal and opportunistic pathogen; and *Candida albicans*, a prevalent component of gut flora, are the top two most common fungal taxa identified on MicrobiomeDB using CORRAL, detected in 364 and 255 samples, respectively.

**Table 1:**
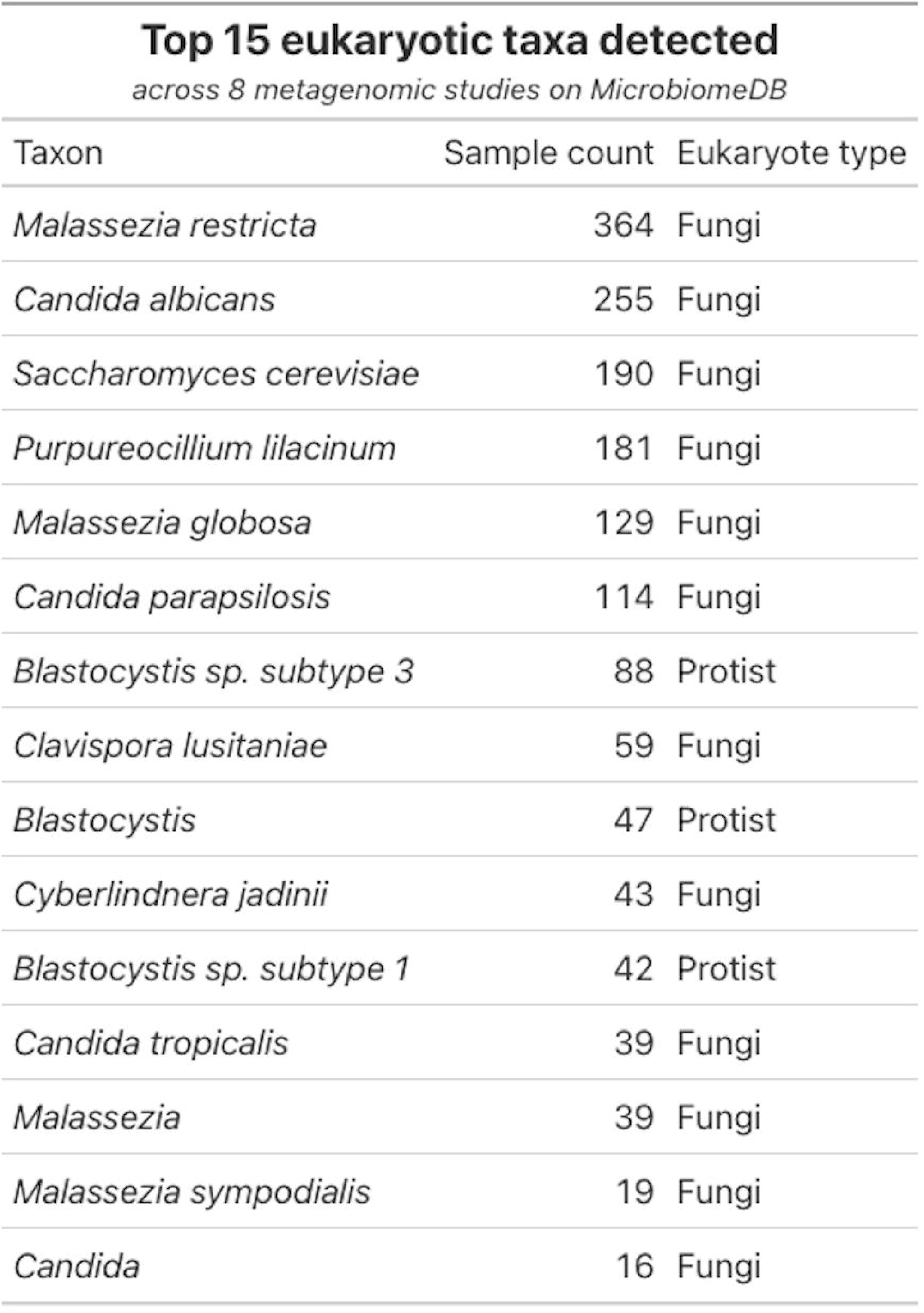
CORRAL expands eukaryote identification when deployed at scale on MicrobiomeDB.org. Top 15 eukaryotes (by prevalence) detected across eight metagenomic studies encompassing 6337 samples.

| Taxon | Sample count | Eukaryote type |
| --- | --- | --- |
| *Malassezia restricta* | 364 | Fungi |
| *Candida albicans* | 255 | Fungi |
| *Saccharomyces cerevisiae* | 190 | Fungi |
| *Purpureocillium lilacinum* | 181 | Fungi |
| *Malassezia globosa* | 129 | Fungi |
| *Candida parapsilosis* | 114 | Fungi |
| *Blastocystis* sp. subtype 3 | 88 | Protist |
| *Clavispora lusitaniae* | 59 | Fungi |
| *Blastocystis* | 47 | Protist |
| *Cyberlindnera jadinii* | 43 | Fungi |
| *Blastocystis* sp. subtype 1 | 42 | Protist |
| *Candida tropicalis* | 39 | Fungi |
| *Malassezia* | 39 | Fungi |
| *Malassezia sympodialis* | 19 | Fungi |
| *Candida* | 16 | Fungi |

### Integration of CORRAL in MicrobiomeDB enables exploration of associations between eukaryotic microbes and host phenotypes

Although CORRAL can be run as stand-alone software, one advantage of integrating this software into MicrobiomeDB is that the results can viewed across many different studies, different sample types, and in many different experimental contexts, thus allowing researchers to identify associations between eukaryotes and study metadata, potentially leading to novel hypotheses (**Figure 6**). The metagenomic data currently available on MicrobiomeDB were generated from distinct geographic regions and from participants that vary in age from pre-term infants, to children, to adults. When we viewed the top 15 most prevalent eukaryotic taxa across all 8 datasets on MicrobiomeDB, in the context of this study metadata, interesting trends emerged. For example, species of *Malassezia* were primarily found in the Human Microbiome Project study (HMP) **(Figure 6A)**, likely because this study included sample types other than stool. A closer look at *Malassezia* species prevalence by sample type across all 8 studies showed that over 60% of the 119 skin and nostril swab samples were positive for *M. globosa*, while *M. restricta* was more restricted to the oral cavity and saliva **(Figure 6B)**. *Blastocystis* sp. were primarily observed in samples from studies carried out in Niger and Malaysia (MORDOR and Malaysia Helminth studies) **(Figure 6A)**, suggesting that these protists may be more prevalent in lower- and middle-income countries. Similarly, *Candida* species were most prevalent in infant samples. The fungi *Clavispora lusitaniae* and *Purpureocillium lilacinum* were each primarily observed in the BONUS-CF and NICU NEC studies, respectively. Interestingly, careful analysis of *P. lilacinum* by the authors of the NICU NEC study identified this organism as a reagent contaminant (31). Taken together, these results suggest that implementing CORRAL at database-scale can accelerate the discovery of species-specific niches, improve the identification of taxa that arise from spurious results or contamination, and help researchers link eukaryotic taxa to environmental covariates within and across studies.

**Figure 6:**
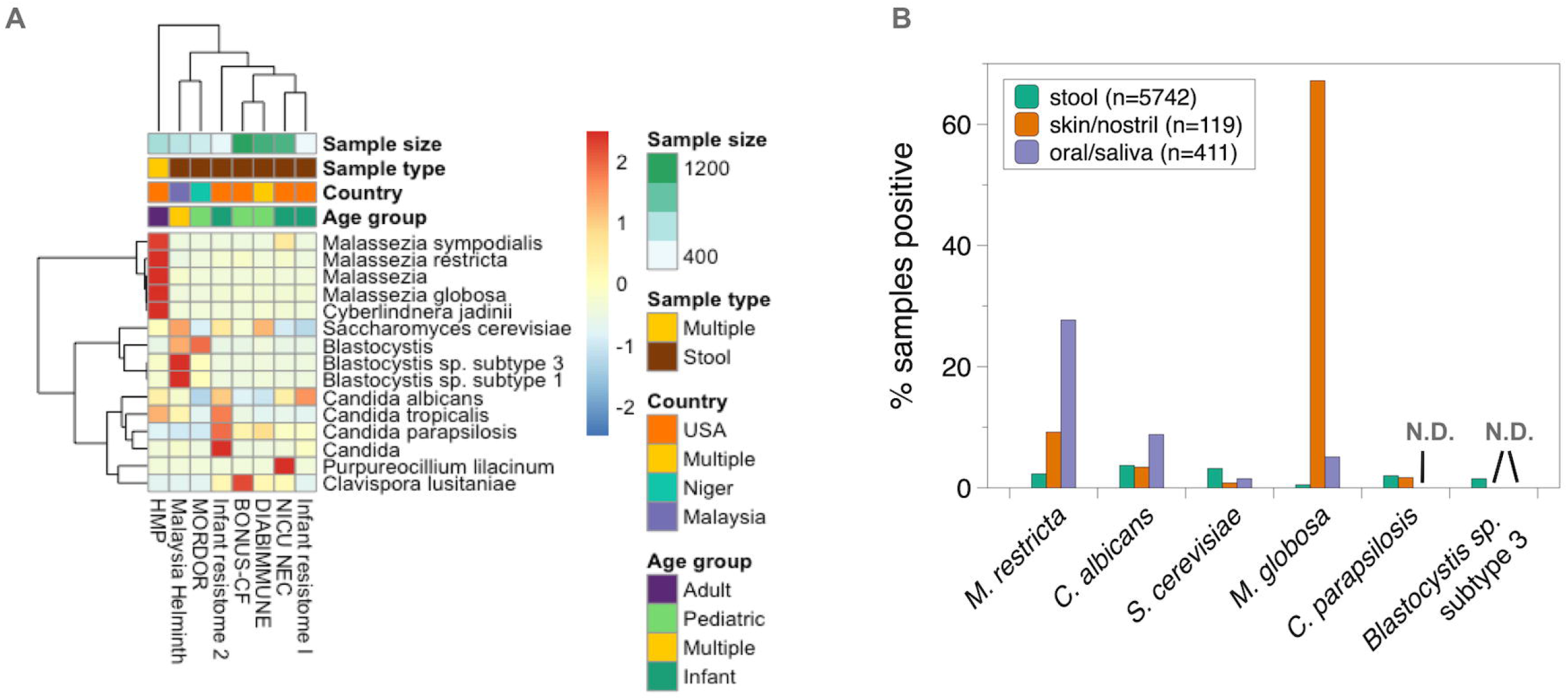
Integration of CORRAL results with study metadata on MicrobiomeDB. A) Heatmap showing row Z scores for the top 15 eukaryotes (by prevalence) across all eight metagenomic datasets currently publicly available on MicrobiomeDB.org. Study name and metadata are shown below and above the heatmap, respectively. B) % of all stool, skin swab or nostril swab (skin/nostril), or oral swab or saliva (oral/saliva) metagenomic samples on MicrobiomeDB.org that were positive for six selected eukaryotes (X-axis) by analysis with CORRAL.

### CORRAL enables quantification of eukaryotes in metagenomic data

In addition to the presence/absence detection of eukaryotes, CORRAL also reports the relative abundance of the eukaryotes it detects, thus opening the door to using many of the same visualization and analytics already familiar to the microbiome community for interpreting bacterial census data. To demonstrate this, we focused on the Human Microbiome Project (HMP) study, since it is the only metagenomic study on our MicrobiomeDB resource that contains multiple sample types. We compared CORRAL’s detection data with relative abundance data for two of the most prevalent fungal taxa detected across all studies on our site, *Candida albicans* and *Malassezia globosa* **(Figure 7)**. Although CORRAL detected *Candida albicans* in less than 10% of vaginal swabs, these positive samples had the highest levels of this organism compared to all other sample types examined **(Figure 7A)**. Although the HMP participants were healthy adults, these data may point to individuals that either had or were at risk of developing vaginal yeast infections. Similarly, *Malassezia globosa* was detected in nearly every skin swab examined **(Figure 7B, left)**, consistent with numerous reports of this fungus as a skin-dwelling microbe, yet the abundance of *M. globosa* is significantly higher in the nasal cavity, compared to skin swabs **(Figure 7B, right)**. These data underscore how quantitative data can impact our understanding of host-microbe interactions. Although this analysis focused on associations between fungal taxa and sample type, a similar analysis could be carried out using any available experimental metadata loaded into MicrobiomeDB (*e*.*g*. correlating fungal taxa with clinical status).

**Figure 7:**
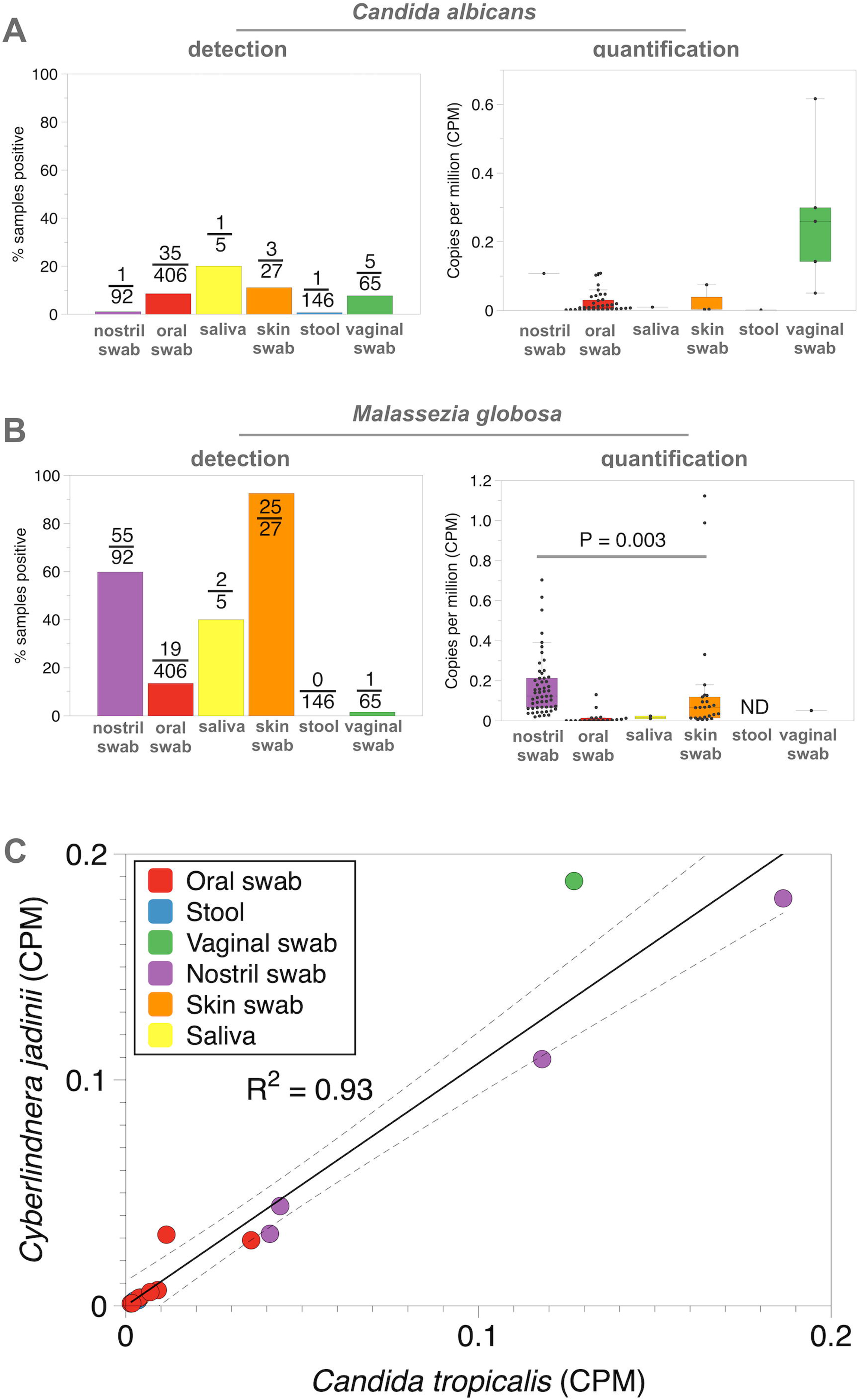
Quantification of eukaryotes by CORRAL. Comparison of detection (presence/absence) and quantification (copies per million; CPM) by CORRAL for A) *Candida albicans* and B) *Malassezia globosa* in the Human Microbiome Project (HMP) study. For detection, number of samples testing positive out of total samples assayed is shown on each bar. P value from Wilcoxon rank-sum test comparing levels of M. globosa between nostril swabs and skin swabs. C) Correlation of CPM for *Candida tropicalis* and *Cyberlindnera jadinii* in HMP.

Quantification data produced by CORRAL also allow conventional statistical analyses to be readily applied, either manually by downloading data from MicrobiomeDB, or directly within the website using data visualization applications (‘apps’) built using the R/Shiny (16,32). For example, we used the ‘Correlation App’ on MicrobiomeDB to search for co-associated fungal taxa. This analysis identified a strong positive correlation between the abundance of the fungi *Candida tropicalis* and *Cyberlindnera jadinii* in the HMP dataset **(Figure 7C;** R^2^ = 0.93). Interestingly, this correlation was evident even in sample types where the relative abundance of these organisms was low or high **(Figure 7C;** oral swabs vs. nostril swabs, respectively). Importantly, due to the relatively low prevalence of eukaryotes in metagenomic samples, observing this type of correlation may only be possible when eukaryotic data can be mined at scale, using large collections of studies. Whether or how these two fungi interact is beyond the scope of this study, nevertheless these data underscore the ability to use CORRAL in conjunction with MicrobiomeDB to generate hypotheses about fungal community interactions which can then be experimentally tested.

## Discussion

CORRAL (Cluster of Related Reference ALignments) is open-source software that uses multiple alignments and Markov clustering to achieve high sensitivity and specificity for identification of eukaryotes in metagenomic data, while also enabling inferences about the presence of eukaryotes not represented in the reference. We highlight the utility of this software using simulated metagenomic samples containing unrepresented species and strains. We also deploy CORRAL on our open-science platform, MicrobiomeDB.org, which allowed automated processing of thousands of samples currently on the site, thus generating the first cross-study atlas of eukaryotes from metagenomic data. With CORRAL now integrated into our standard data loading workflow for metagenomic studies on MicrobiomeDB, this atlas will continue to grow as new studies are loaded. This demonstrates the value of combining robust software with web-based tools for conducting large-scale screens of metagenomic data, thereby creating a resource that will allow investigators to access eukaryotic data from a vast range of sample types and studies, irrespective of whether the original study investigators intended to examine eukaryotes in their data.

The high cost of metagenomic sequencing, the relative low abundance of most eukaryotes in the microbiome, and the inherent limitation of reference-based methods for identification of taxa remain major challenges to identification of eukaryotes. CORRAL helps to address some of these issues by being able to work with minimal information required to plausibly report the presence and abundance of eukaryotes, even when the source reads do not perfectly match the marker gene reference. Future improvements in genome assembly will provide more complete information on eukaryote-specific genomic sequences which could be used to create a larger reference with more taxa and more sequences per taxon, improving both specificity and sensitivity of hits reported by CORRAL.

It remains to be seen how well CORRAL, or for that matter any eukaryotic prediction tool, compares with standard diagnostics used to identify microbial eukaryotes in biological samples. Microscopic examination of stool is one of the most common methods for detecting and diagnosing common infections with protozoan parasites, helminths, and fungi, but these methods require significant enrichment, for example by Baermann float or sucrose centrifugation, or involve staining with special dyes to allow detection even when microbes are present at extremely low abundance. In addition, microbial eukaryotes are often highly resistant to common lysis conditions used for DNA extraction prior to metagenomic sequencing (e.g., fungal spores or *Cryptosporidium* oocysts), which means that they are likely to be poorly represented in DNA preparations used for metagenomics. Since curated sample metadata is loaded together with microbiome data for each study represented on MicrobiomeDB.org, we can explore the relationship between diagnostic assay results and metagenomic results. For example, the Malaysia Helminth metagenomic study currently available on MicrobiomeDB.org specifically addresses the impact of helminth infection on the microbiome and thus includes sample metadata for different helminth species detected in 650 stool samples from over 400 participants. Nearly half of all samples tested positive by microscopy for either *Ascaris*, hookworms, or *Trichuris*, yet none of these organisms were detected in any samples by either EukDetect or CORRAL (data not shown; MicrobiomeDB.org). Based on these discrepancies, we anticipate metagenomic sample preparation – and consequently any downstream analysis with tools like CORRAL – likely results in an under-estimation of eukaryotic microbes. For this reason, we do not recommend using outputs from CORRAL (or other available detection tools for microbial eukaryotes) in downstream alpha or beta diversity analyses. Even with this potential under-estimation, *Candida albicans* and *Candida parapsilosis* were the second and sixth most common eukaryotes detected by CORRAL on MicrobiomeDB, and were recently designated as critical or high priority group fungal pathogens by the World Health Organization (33).

Our data suggest that the ability of any software tool to distinguish related eukaryotic taxa may depend on multiple factors including the exact species in question **(Figure 1)**, sequence divergence of an organism from the reference **(Figure 2)**, the presence of closely related species in the same sample **(Figure 5)**, and the sequencing depth and/or abundance of these organisms in the sample **(Figure 4A and 5)**. Therefore, the selection of software should, as always, be guided by the specific question and goal of the analysis.

Our strategy of clustering of related read alignments could be further improved by making use of information about taxonomic similarity between reference sequences. Not relying on external data about similarity of different proteins has the benefit of flexibility but lacks the capacity to act on implied ‘improbability’ of reported taxa. For example, it is relatively unlikely that a sequenced sample containing reads which map to multiple closely related *Leishmania* species does in fact contain different species of *Leishmania*, because the reference sequences are highly similar, and the species readily hybridize [31]. Conversely, reads sharing alignments to markers across a large taxonomic distance are more likely to come from a single source because of relative implausibility of the sample containing multiple eukaryotes of unknown genera – for example, they might all be contamination from a metazoan host. Incorporating such speculations about ‘likely’ and ‘unlikely’ results into a detection method is an ambitious undertaking, because it involves making and modeling assumptions about vast numbers of eukaryotic taxa, most of which have not been sequenced and not yet well studied. It could, however, yield methods with a more natural choice of threshold parameters, and further gains in sensitivity and specificity. Finally, since the computational approach used by CORRAL is independent of the reference sequences used, our software could potentially be applied to processing alignments to any reference that is anticipated to be redundant and incomplete, and where reads are expected to map with varying identity. This includes identification of bacteria to the strain-level resolution required in genomic epidemiology, as well as taxonomic classification of viral reads to reference sequences (34), identification of antibiotic resistance genes (35), or bacterial virulence genes (36).

### Conclusion

CORRAL enables detection of eukaryotic organisms in shotgun metagenomics data at scale. We build on the strategy of aligning data to a previously curated set of conserved eukaryotic genes and employ an approach to incorporating alignment evidence based on using multi-aligned reads for MCL clustering, which results in an increased sensitivity, improved quantification, and reporting of uncertainty in predictions. CORRAL has been integrated in MicrobiomeDB.org, creating a publicly accessible cross-study atlas of eukaryotic microbial content in public datasets.

### Implementation

#### CORRAL

CORRAL is implemented as a Nextflow workflow that downloads raw sequence files, produces alignments to a reference database using Bowtie2, and processes these alignments using our custom Python package, `marker_alignments`. This package uses the Python module `pysam` to read alignments into an SQLite table where each row corresponds to one alignment, storing information about: the identifier for each read; name of matched marker and taxon; contribution to coverage (fraction of marker covered by the alignment), and match identity (fraction of bases agreeing between query and reference). Counts of entries in this table, along with the coverage field are used for quantification, and the identity field is used for clustering and filtering. We then apply filters and construct a marker similarity graph with SQL queries, run MCL using a `markov_clustering` package for marker and taxon clustering, and group the taxa by taxon cluster. We then apply another SQL query to produce quantified results. The main challenge in developing the software was arriving at an adequate understanding of how alignments to markers may look when the source taxon is not the same as the reference taxon, then capturing this information as a procedure that yielded sensible results. We addressed this challenge by organizing the software as a sequence of parametrizable filters and transforms, which allowed us to evaluate candidate procedures on a wide range of empirical data. This flexibility is built into the software, which allows any user to override the default behavior of CORRAL.

#### Simulations and comparisons

Software used to prepare data for this publication was written as a mix of reference tools with custom Python and Bash tools. These tools simulated data, ran EukDetect and CORRAL pipelines, cross-checked with databases, prepared spreadsheets available to the reader as supplemental material, and more. To overcome the challenge posed by the complexity of the process, we wrote developed the tools iteratively while generating evidence, tracked changes with Git, and organized them into a `Make` pipeline, which helped us build on previously generated results during iterations and keep track of our work.

### Availability and Requirements

Project name: CORRAL

Project home page: github.com/wbazant/CORRAL

Operating system(s): POSIX compatible system (Linux, OS X, *etc*.).

Programming language: Nextflow, Python

Other requirements: Bash 3.2 or higher, Java 11 or higher

License: MIT

Any restrictions to use by non-academics: none

## Methods

### CORRAL workflow

CORRAL is a Nextflow pipeline wrapping a Python module we created, called marker_alignments, and combining it with existing tools for retrieving and aligning raw reads. Marker_alignments is installed using the package manager Pip (pip install maker_alignments). This Python package carries out all Markov clustering steps outlined in **Figure 3** and is easily parameterized by users.

CORRAL uses an ≥ 97% match identity threshold for unambiguous matches. Clusters with an average match identity < 97% but that have at least 4 reads aligning to 8 markers are designated as ‘strong’ ambiguous hits, possibly indicating a taxon that is not represented in the reference marker database but which is related to taxa that are in the reference. Ambiguous hits that have fewer than 4 reads aligning to 8 markers are considered weak evidence and are not reported by CORRAL. The 3% threshold was selected empirically based on evaluating samples with ≥ 90% identity for read mapping, inspecting these read alignments at the level of individual markers, and manually classifying them as either preferentially aligning to one reference species, or aligning to multiple reference species. Also, by default, CORRAL rejects taxa for which ≥ 50% of the marker genes in the cluster fall below the cluster average **(Figure 3, step 4)**. We relate the values to the marker average per cluster since when one taxon shows higher quality alignments than the others, we want to count it as evidence towards the taxon being present, regardless of the absolute strength of the match. We also want inferior alignments in a marker cluster to count as evidence against a taxon being present (a more plausible explanation for lower quality alignment is that the taxon is a different species) and the 50% threshold corresponds to balancing evidence from different clusters by treating them as independent and equally valuable pieces of information. All threshold values in CORRAL are overridable, allowing users to modify default settings to achieve different outcomes (e.g., lowering a threshold to increase sensitivity).

CORRAL quantifies abundance for each found taxon with ‘copies per million’ (CPMs) as the number of reads assigned to the taxon normalized by marker length and sequencing depth, in line with the quantity calculated by the integrated metagenomic profiling tool, HUMAnN (37).

### Simulations and mock community analysis

For all simulations, wgsim (38) was used to sample 100 basepair reads with base error rate of 0 from the EukDetect reference (the 1/23/2021 version, latest at time of writing, consisting of BUSCOs from OrthoDB (39)). Bowtie2 (17) was then used to align reads back to the references with identical settings to those used in EukDetect: the end to end (default) mode and the `--no-discordant` flag. To assess the correctness of simulated alignments, we retrieved the rank of the nearest taxon containing source and match by using the ETE toolkit and the NCBI database version dated 2020/1/14 packaged with EukDetect. Alignments were deemed correct if the source and match were of the same species, or genus in case of hold-out analysis where the species was missing from the reference by design. Precision and recall were calculated using the OPAL method of assessing taxonomic metagenome profilers (40), where precision is a fraction of correctly mapped reads among all reads that are mapped, and recall is a fraction of correctly mapped reads among all reads. When simulating whole samples, we obtained 338 simulated samples from a holdout set of 371 taxa, because we skipped 33 cases in which `wgsim` considers the sequences too fragmented to source reads at a set coverage, and errors out. The number of reads sourced per marker to obtain 0.1 coverage was calculated as previously described (41).

For analysis of closely related pairs of species, we subsampled our dataset of 4558 data points using computed PCA coordinates. 863 pairs where one or both members were not reported at the species level were excluded, and the remaining 3695 pairs were passed to a greedy geospatial subsampling algorithm with the distance parameter 0.005 and a seed pair of *Entamoeba hystolytica* and *Entamoeba dispar* used in the original EukDetect publication. This yielded 50 data points, constituting a “focused subset” of low complexity pairs falling close to the *E. hystolytica* and *E. dispar* pair (Figure 5D-I). To generate a “broad subset” that would be more skewed toward high confusability, we reran this algorithm with the distance parameter set to 0.05, resulting in 98 data points **(Figure 5J-O)**.

Mock microbial community data was obtained from the ZymoBIOMICS Microbial Community Standard (catalog no., D6300) and was retrieved from the Sequence Read Archive (PRJEB38036) and processed using either the standard EukDetect or CORRAL workflows. Alternative analyses were also carried out in which Bowtie2 was used to simply map all reads from each mock community sample to the EukDetect marker gene reference, filtering out alignments that were less than 60 bp, keeping only the best alignment for each marker gene, and either including or excluding a MAPQ filter **(Figure 4C, 4D, and Table S1)**.

### Deploying CORRAL on MicrobiomeDB.org

CORRAL is integrated into the standard MicrobiomeDB workflow for metagenomic datasets (see *https://github.com/VEuPathDB/MicrobiomeWorkflow*) along with bioBakery tools for bacterial abundance estimation. CORRAL output results areis loaded as both binary (presence/absence) and quantitative Copies Per Million (CPM) values for each sample and can be used along with other sample details related to the collection, processing, and analysis of data for filtering and stratification of bacterial abundance data as well as directly for exploring correlations between eukaryotic abundance and other sample data. Strong ambiguous results from CORRAL **(Figure 3, step 7)** are reported on MicrobiomeDB at the genus level with the ‘?’ removed.

## Declarations

### Ethics approval and consent to participate

No applicable for this study

### Consent for publication

All authors have reviewed this manuscript and consent to its publication

### Availability of data and materials

All our software is freely and publicly available under the MIT license: CORRAL (*github*.*com/wbazant/CORRAL*), its main Python module, (*github*.*com/wbazant/marker_alignments*). Fully reproducible code associated with all simulations, greedy geospatial subsampling, and other analyses in this manuscript (including Python, Make, and Bash scripts) are available on Github (*github*.*com/wbazant/markerAlignmentsPaper*). All CORRAL results are publicly viewable and downloadable on the open-science resource, MicrobiomeDB.org.

### Authors’ contributions

D.P.B., W.B., and KC designed the experiments and wrote the paper. W.B. wrote all code and carried out simulations and evaluations of CORRAL and EukDetect. D.P.B., W.B., and KC designed figures. A.B. clarified the description of the CORRAL method, illustrated the main schematic for CORRAL, and provided feedback and edits on manuscript drafts.

## Acknowledgements

We thank the authors of the metagenomic studies cited in this work (22–29) for their assistance with loading and representing their data on MicrobiomeDB.org.

## Figure legends

**Supplementary Table 1: Analysis of mock community standard**. Table showing alignment results for 11 different methods applied to ZymoBIOMICS mock community standard (D6300). A portion of this data was used to generate Figure 4C and 4D. Raw data for this analysis was obtained from Sequence Read Archive accessionn PRJEB38036 and is described in Yang et al. (21).

**Supplementary Table 2: Confusability analysis for pairs of eukaryotic microbes**. Table showing all results for confusability analysis described in Figure 5.

**Supplementary Figure 1: Mathematical framework for evaluating the potential confusability of closely related pairs of eukaryotic species present in the same sample**. (A) Schematic showing all eight parameters of the model for relating read mapping within and outside of a given pair of taxa. These eight parameters were calculated for all 4558 pairs of taxa. (B) Table showing four example pairs of taxa, each with different read mapping behavior. (C) Principal component analysis (PCA) plot showing first two principal components from the analysis of the full dataset from panel B. Axes show interpretation of principal components. Color points represent the position of example taxon pairs from panel B.

**Supplementary Figure 2: CORRAL results are readily searchable and queryable on MicrobiomeDB.org**. Screenshots showing CORRAL results for the DIABIMMUNE study that includes 1149 samples. Results are displayed as detection (top) or quantification (bottom). Users select broad groups of microbial eukaryotes (e.g. (A) fungi) and the select specifc species (B) from a multipick list. Numbers of samples in which the taxon was detected are shown and can be used to filter the dataset (C). For quantification, users select the ‘normalized’ number of sequence matches (D) and then a species of interest (E) to display a histogram of sample count by abundance for the the taxon of interest, making it simple to identify samples with high (F) or low abundance. Users can then select only those samples that contain a specific abundance or fall within an abundance range.

