## Supplementary figures and images for "Improved eukaryotic detection compatible with large-scale automated analysis of metagenomes"

### Figure S1

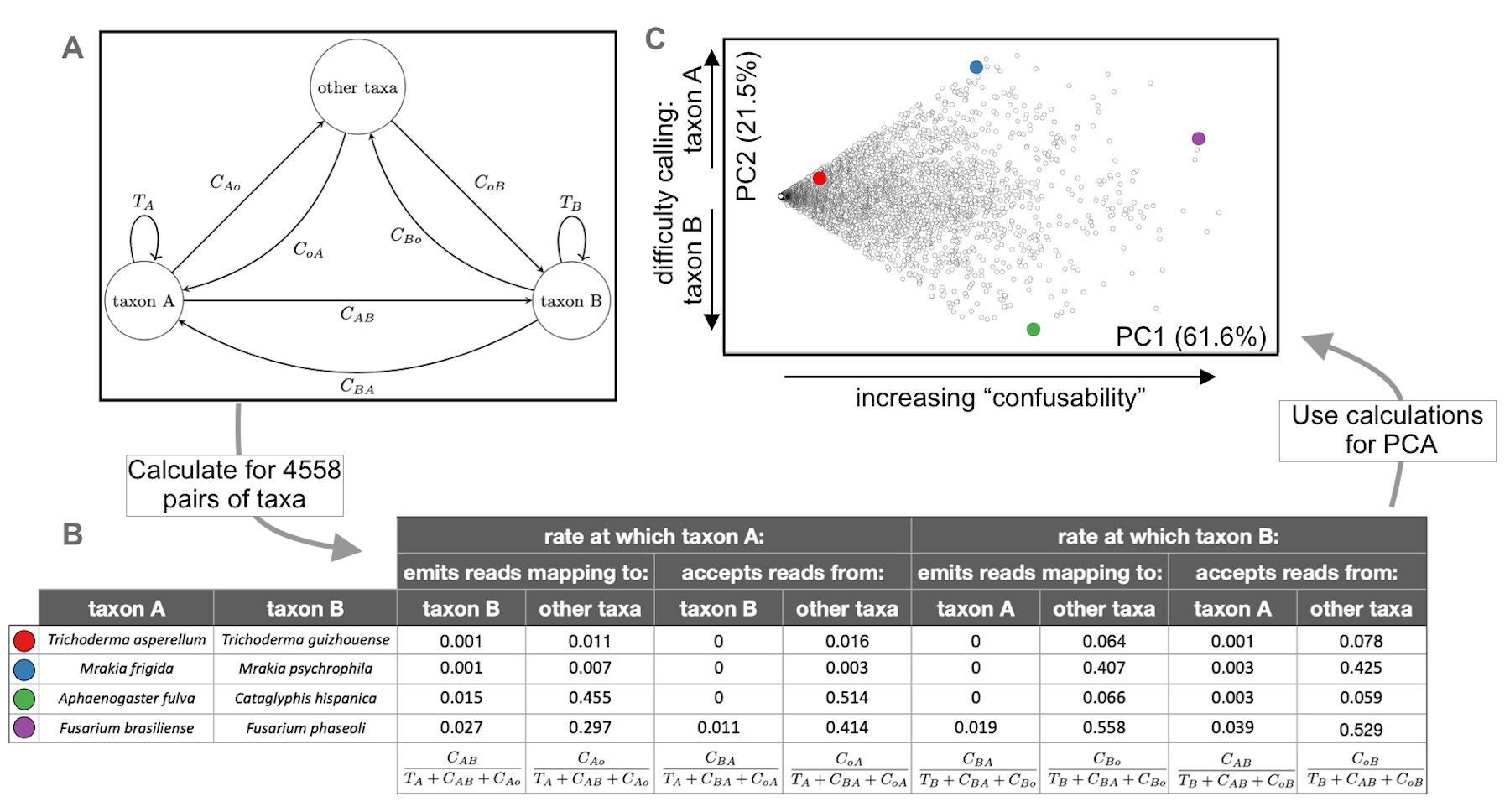

### Figure S2

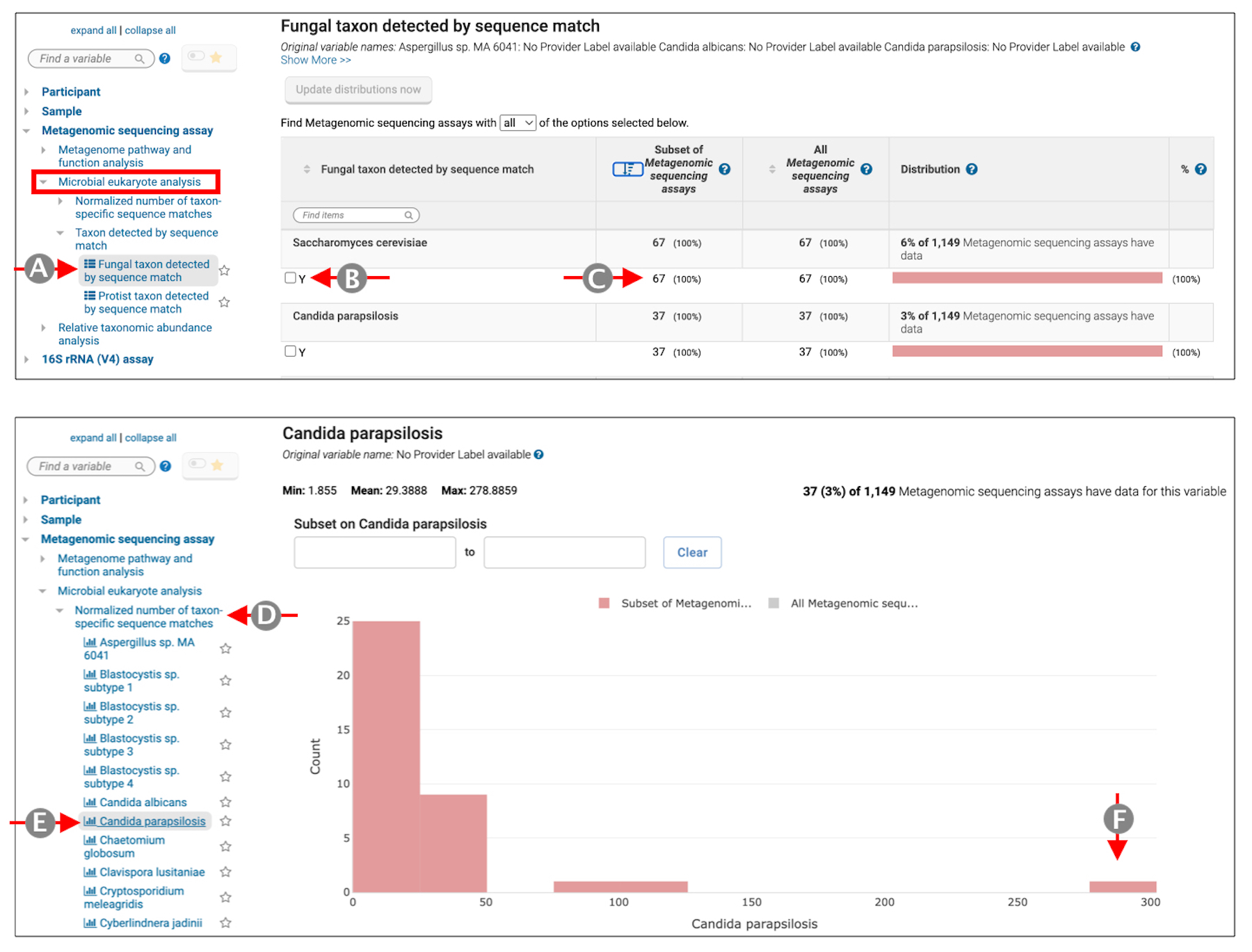
